# Multicellular nodules form at subacute time points following closed-head diffuse traumatic brain injury in pigs

**DOI:** 10.64898/2026.11.24.768113

**Authors:** Susan Shin, Adrina Tiju, Caela C. Long, Nylah Miles, Ahmed Altaf, Kevin D. Browne, Mariko L. Bennett, D. Kacy Cullen, David F. Meaney, Kathryn L. Wofford

**Author notes:** Corresponding author **Correspondence and reprint request should be directed to**: Dr. Kathryn L. Wofford, Department of Neurosurgery, University of Pennsylvania, 105 Hayden Hall, 3320 Smith Walk, Philadelphia, PA, 19104, USA.

## Abstract

Traumatic brain injury (TBI) is a leading cause of long-term neurological impairments and is associated with persistent neuroinflammation. Multicellular nodules, composed of densely aggregated myeloid cells, have been reported in human post-mortem tissue following TBI but remain poorly characterized. Here, we utilized a clinically relevant porcine model of closed-head diffuse TBI generated through rotational acceleration to characterize the temporal and spatial distribution of these nodules. Nodules were rare in sham animals and minimally present at acute time points but increased significantly by 14 days post injury. Nodule burden increased as a function of injury severity and was positively correlated with rotational injury velocity. Nodules were enriched within white matter structures, including subcortical white matter, corpus callosum, fornix, and internal capsule, while remaining minimally present in cortical gray matter. Within these regions, nodules exhibited a widespread yet spatially organized distribution rather than local clustering around a single anatomical location. Together, these findings demonstrate that multicellular nodules are a robust and reproducible neuroinflammatory pathology following diffuse TBI in a gyrencephalic large-animal model. By establishing the temporal progression, severity dependence, and white matter enrichment of this clinically relevant but understudied endophenotype, this work provides a foundation for future investigations into the mechanisms and consequences of post-traumatic neuroinflammation.

## INTRODUCTION

Traumatic brain injury (TBI) is a major public health concern, affecting approximately sixty-nine million people globally each year ^1^. TBI is associated with persistent long-term neurological impairments that may persist for months to years after the initial injury, including deficits in memory, attention, processing speed, and executive function ^2–8^. While the initial injury response to TBI is adaptive, secondary injury cascades, such as inflammation and microglial activation, can contribute to long-term impairments ^9^.

Microglia, the brain’s resident parenchymal macrophages, survey the healthy brain to maintain homeostasis and respond to signs of damage or disease. Following a TBI, microglia respond within minutes by migrating to the injury site, proliferating, and undergoing morphological transformation to clear cellular debris and initiate repair ^10–16^. This early activation is largely neuroprotective, as microglia respond to injury and release cytokines and neurotrophic factors that support tissue integrity and neurological recovery ^17,18^. However, if microglia remain persistently activated, they can hinder CNS repair and contribute to secondary neurodegeneration ^17–20^.

Microglial morphology is one way researchers have examined the response of microglia following TBI in preclinical and clinical studies ^21–24^. Under homeostatic conditions, microglia often take on a ramified morphology and upon sensing damage or infection, microglia can change their morphology to hyper-ramified, amoeboid, rod-shaped, or dystrophic ^25–28^. Beyond these common and well characterized single-cell morphologies, microglia have also been reported to aggregate into nodules–multicellular clusters of densely packed cells. This nodule morphology (also termed stars, clusters, clumps, knots, and aggregates in the literature) has been observed in human post-mortem tissue samples collected sub-acutely and chronically after TBI ^29–35^ and in other disease contexts such as multiple sclerosis and encephalitis ^36–39^. Despite their clinical relevance, nodules remain poorly characterized, in part because pre-clinical TBI models rarely generate robust nodule development.

Here, we report that a clinically relevant model of closed-head rotational acceleration diffuse TBI in pigs promotes abundant nodule development. To better understand this TBI-induced endophenotype, we investigated nodule burden, density, and location across time, brain region, and injury severity. Characterization of nodule development and distribution is the first step in understanding this clinically relevant and understudied TBI-induced immune response. This work demonstrates that this preclinical model can reliably and robustly generate this understudied endophenotype, allowing future investigations into the stimuli, features, contributions, and consequences of TBI-induced nodules.

## METHODS

### Animal enrollment and handling

Animal housing and care for animals utilized within these studies were in accordance with the Guide for the Care and Use of Laboratory Animals ^40^ and followed ARRIVE Guidelines ^41^. Procedures received prior approval from the University of Pennsylvania’s Institutional Animal Care and Use Committee. 6-month-old adult female Yucatan minipigs (average weight of 31.3 kg) acquired from Sinclair Research were utilized. Upon arrival, animals underwent a standard quarantine period to confirm an absence of transmissible zoonotic pathogens and to facilitate environmental acclimation. Animals were housed indoors in a facility accredited by the Association for Assessment and Accreditation of Laboratory Animal Care International with water available *ad libitum*.

Animals were randomly assigned to a sham procedure (n=3), a moderate angular velocity injury procedure (n=4), or a high angular velocity injury procedure (n=7). Animals in the high angular velocity group were survived for either 3 (n=3) or 14 (n=4) days post-injury (dpi). All subjects in the sham and moderate angular velocity injury groups were survived to 14 dpi. Animals were fasted 12 hours prior to procedure with water remaining available *ad libitum*.

Animals were induced via an intramuscular injection combining ketamine (20–30 mg/kg) and midazolam (0.4–0.6 mg/kg). Following initial delivery of 5% isoflurane via a snout mask, subjects were intubated with a size-appropriate endotracheal tube alongside administration of subcutaneous glycopyrrolate (0.01 mg/kg). Anesthesia was subsequently maintained using isoflurane (titrated within a 1–5% range, typically 1.5–2.0%) delivered in 2–3 L of pure oxygen per minute and adjusted dynamically to maintain targeted physiological parameters: heart rate between 100–130 beats per minute, respiratory rate of 9–12 breaths per minute, and arterial oxygen saturation ranging from 97% to 100%. Core body temperature was actively stabilized throughout the procedure using a forced-air warming unit and thermal blankets. Protective lubricating gel was administered on the eyes to prevent corneal drying.

### Closed-head diffuse brain injury

To induce a closed-head, diffuse rotational acceleration injury, anesthetized animals were secured to a padded bite plate, connected to a HYGE pneumatic actuator. Adjustable snout cables were used to carefully position and anchor the mouth to the apparatus. The HYGE device delivers purely impulsive, non-impact rotational movement, which produces rotational forces scalable to clinical TBIs ^42–44^. In this study, animals underwent sagittal plane rotation (aligned with the brain stem) at two levels: a moderate angular velocity injury (average 88.5 radians per second (rad/s)) or a high angular velocity injury (average 107.0 rad/s). Angular displacement over time was recorded in LabVIEW with two magneto-hydrodynamic sensors (Applied Technology Associates, Albuquerque, NM) operating at a sampling rate of 10 kHz. Angular velocity was post-processed following SAE J211 standards using Butterworth filter with a cutoff frequency of 300 Hz in MATLAB R2024b (MathWorks – Natick, MA), as previously described [45]. Sham animals received all handling and anesthetic procedures excluding the rotational acceleration. Injuries generated at these kinematic levels are scalable to human data and approximate a less injurious clinical TBI (previously termed mild) and a moderately injurious clinical TBI for moderate-velocity and high-velocity TBI groups, respectively ^45^.

Upon completion of the rotational injury procedure, animals were uncoupled from the bite plate, inspected for oral or dental injuries, and transported back to their housing units. Isoflurane delivery was terminated and a sustained-release formula of buprenorphine (0.1 mg/kg) was administered subcutaneously for analgesia. Endotracheal extubation was performed once subjects exhibited protective reflexes such as chewing, swallowing, or coughing. Post-operative status was tracked until animals were ambulatory and stable and daily thereafter.

### Terminal procedure and tissue collection

At the respective experimental endpoints, animals were weighed, anesthetized, and intubated as described above. Subjects underwent transcardial perfusion using 0.9% heparinized saline, followed by 10% neutral buffered formalin. The head was removed and placed in formalin for an initial 24-hour post-fixation period. Brain tissue was then extracted and stored in fixative for an additional week at 4°C. We observed no gross pathological changes or procedural effects such as contusions or lesions in all animals, consistent with our previous work ^13,43,46,47^.

### Brain tissue histology

Coronal blocking of brains began at the optic chiasm, producing 5 mm thick tissue blocks. These blocks were processed and paraffin embedded, after which 8 µm sections were sectioned using a manual rotary microtome. Chromogenic 3,3’-Diaminobenzidine (DAB) immunohistochemistry was performed following manufacturer’s protocol (Vector Laboratories DAB Substrate Kit, Vector Labs, SK-4100). Briefly, slides underwent tissue rehydration, heat-mediated antigen retrieval via a pressure cooker in Tris-EDTA buffer (pH 8.0), blocking with normal horse serum, overnight incubation in primary antibody, and subsequent signal visualization (VECTASTAIN Elite ABC Kit, Vector Labs, PK-6200). Immunostaining targeted ionized calcium-binding adapter molecule 1 (Iba1; Wako, 019-19741; 1:4000 dilution), paired with a hematoxylin counterstain (Hematoxylin solution Gill No. 3; Sigma-Aldrich, GHS332-1L). To minimize batch variability, all tissue sections were processed simultaneously within the same histological run.

### Imaging

Images of DAB-stained sections were captured at 20X magnification using an Aperio CS2 slide scanner (Leica Biosystems Inc., Buffalo Grove, IL). Given the possible morphological variations of microglia across distinct anatomical zones, whole brain images were utilized to capture comprehensive Iba1 expression profiles for all animal subjects.

### Pathological characterization: Regions of Interest definition

QuPath (version 0.7.0), an open-source digital pathology platform, was utilized to analyze images. Each brain section was divided into four anatomical regions of interest (ROIs): ROI 1 consisted of cortical gray matter; ROI 2 included subcortical white matter, corpus callosum, and claustrum; ROI 3 contained hippocampal formation and fornix; and ROI 4 encompassed basal ganglia, thalamus, and internal capsule. Each ROI was manually selected on whole brain images.

To compare nodule burden and density between gray matter and white matter regions in the 14 dpi high-velocity injury group, we defined gray matter nodules as those within ROI 1 and white matter nodules as those located exclusively within white matter structures of ROI 2 (corpus callosum and subcortical white matter) and ROI 3 (fornix). ROI 4 was omitted from this analysis because tissue boundaries made it difficult to reliably differentiate gray matter from white matter.

### Automated detection of nodules

An automated detection pipeline was developed in QuPath to identify potential Iba1+ nodules within each ROI. To optimize processing, each ROI was subdivided into 5000 x 5000 μm tiles, and a custom script iteratively analyzed selected tiles until the entire ROI was covered. Within each tile, potential nodule objects were detected using a custom pixel classifier generated via the “Create thresholder” workflow. Images were processed at full resolution of 1.87 μm/pixel in the DAB channel using a pixel threshold of 0.7 without a smoothing prefilter. Objects larger than 230 μm² were annotated as potential nodules. This area threshold was determined to exclude some minor tissue folds and all singlet microglia while including the smallest nodules identified on manual review.

### Manual nodule classification and quantification

Following automated detection, all potential nodules were manually reviewed using the nodule classification process and false positives were excluded. Selected objects that passed manual verification and were considered definite nodules were categorized, quantified, and reported here. Detected potential nodules were first visually screened at 1X magnification based on their appearance as large, densely stained clusters of Iba1+ immune cells distinct from the surrounding tissue. Artifacts from procedural debris, tissue folds, or background staining were removed. Then, potential nodules were inspected at 7X magnification to verify they met nodule criteria, which was defined as a compact, multicellular cluster of Iba1+ cells forming a globular shape exceeding an area of 230 μm², distinct from surrounding microglia. Nodule burden was quantified as the total count of validated nodules within the area of interest, and nodule density was calculated by dividing total nodule count by the tissue area (mm²).

### Statistical analyses

Comparisons of the unpaired variables across survival time point groups and injury severity groups were evaluated using ordinary one-way analyses of variance (ANOVAs) followed by Tukey’s post-hoc tests corrected for multiple comparisons. Regional comparisons across ROIs were assessed using repeated measures one-way ANOVAs with Geisser-Greenhouse corrections and Tukey’s post-hoc tests. Comparison between white matter and gray matter nodules was assessed through paired t-tests. A linear regression was completed using ordinary least squares estimation. All statistical tests were completed in GraphPad Prism version 10.4.1 (532).

## RESULTS

### Myeloid cells formed nodules in the pig brain following TBI

Following TBI, we observed ionized calcium-binding adapter molecule 1-positive (Iba1+) immune cells, presumed to be predominantly microglia, formed numerous, dense, multicellular aggregates or clusters throughout the brain (Fig. 1A-C). These aggregates, rarely present in sham tissue, were most notable in animals surviving 14 days post high-velocity TBI. These formations, herein termed nodules, were distributed throughout the brain - detectable in multiple anatomical regions rather than concentrated at a single site. Furthermore, ramified microglia typically spanned the spaces between nodules, allowing their formation to be easily detectable.

**Figure 1:**
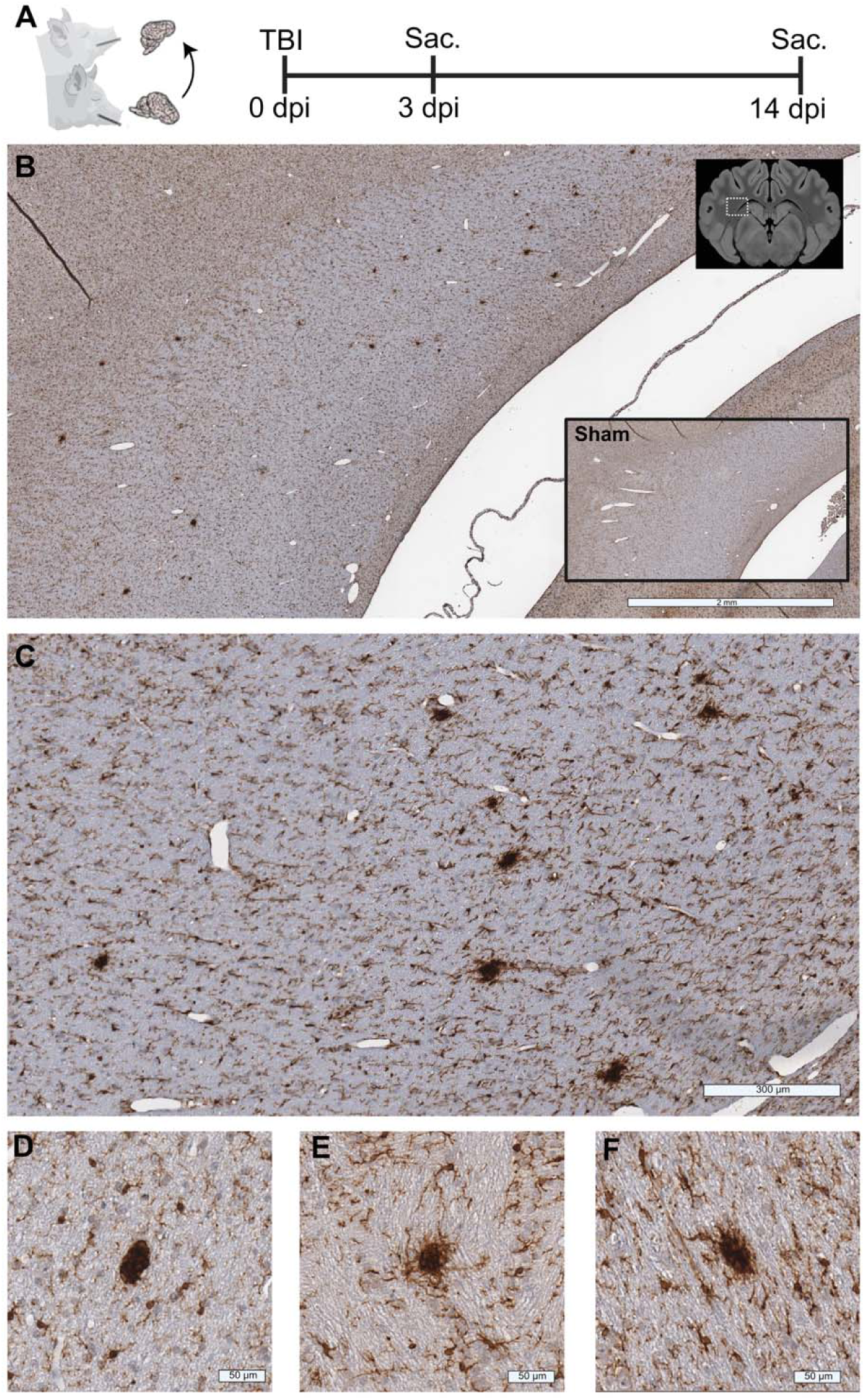
Nodules are abundant following closed-head rotational TBI in pigs. **(A)** Schematic of the closed-head rotational acceleration TBI model in pigs and the associated timeline indicating the injury or sham procedure as well as the terminal sacrifice (Sac.) time point. (**B)** Representative low-magnification image of Iba1 immunohistochemistry (IHC) demonstrating nodules within the brain following TBI. The upper inset indicates the approximate anatomical region represented in the histological image. A representative sham section is shown in the lower-left inset for comparison (scale bar = 2 mm). (**C)** Higher-magnification of the injured tissue with multiple discrete, densely stained Iba1+ nodules distributed throughout the tissue (scale bar = 300 μm). (**D–F)** Representative high-magnification images of injured tissue demonstrating the different morphologies of microglia nodules (scale bars = 50 μm).

Given these qualitative observations, we sought to define and quantify nodule burden and density across injury severity, time, and region. Nodules were defined as a compact, globular group of Iba1^+^ cells greater than 230 μm² with strong Iba1 staining, exhibiting distinct morphology relative to neighboring cells. At low magnification, nodules appeared highly homogeneous in size and shape. When viewed at high magnification, nodule size, shape, and ramifications exhibited some variation but still allowed easy discrimination from neighboring ramified microglia (Fig. 1D-F).

### Nodule burden increased at subacute time points following TBI

To investigate the temporal relationship of nodule formation following TBI, we analyzed a cohort of pigs that experienced a sham or high-velocity TBI and survived to an acute time point (3 dpi) or a subacute time point (14 dpi). In sham animals, most Iba1^+^ cells were evenly spread throughout the tissue and exhibited a ramified morphology visible at low and high magnifications (Fig. 2A-C). Rare nodules were observed in sham animals with 12 4 (mean SEM) nodules throughout an entire coronal cross-section of brain tissue (Fig. 2D). At the acute (3 dpi) time point, nodules were minimally present with most of the Iba1^+^ cells evenly distributed throughout the tissue (Fig. 2D). There was a non-significant increase in the mean number of nodules at 3 dpi (55 15) throughout the entire brain section compared to sham animals (Fig. 2D; p=0.7222). In contrast, at 14 dpi, nodules were abundant within the tissue (Fig. 2A-B). At 14 dpi, there was a mean 271 49 nodules through the brain section and nodule burden at 14 dpi was significantly greater than sham (p=0.0033) and 3 dpi (p=0.0087; Fig. 2D). Small differences in neuroanatomy and blocking procedures can alter the tissue cross-section. To account for changes in tissue area across animals, we normalized nodule burden to tissue area. The robust temporal trend in nodule burden was maintained after area normalization, with nodule density at 14 dpi significantly greater than both sham animals (p=0.0087) and the acute time point (3 dpi) group (p=0.0175; Fig. 2E). Thus, this trend suggests the accumulation of nodules reflects a pathological progression dependent on time with abundant nodule numbers developing at subacute time points.

**Figure 2:**
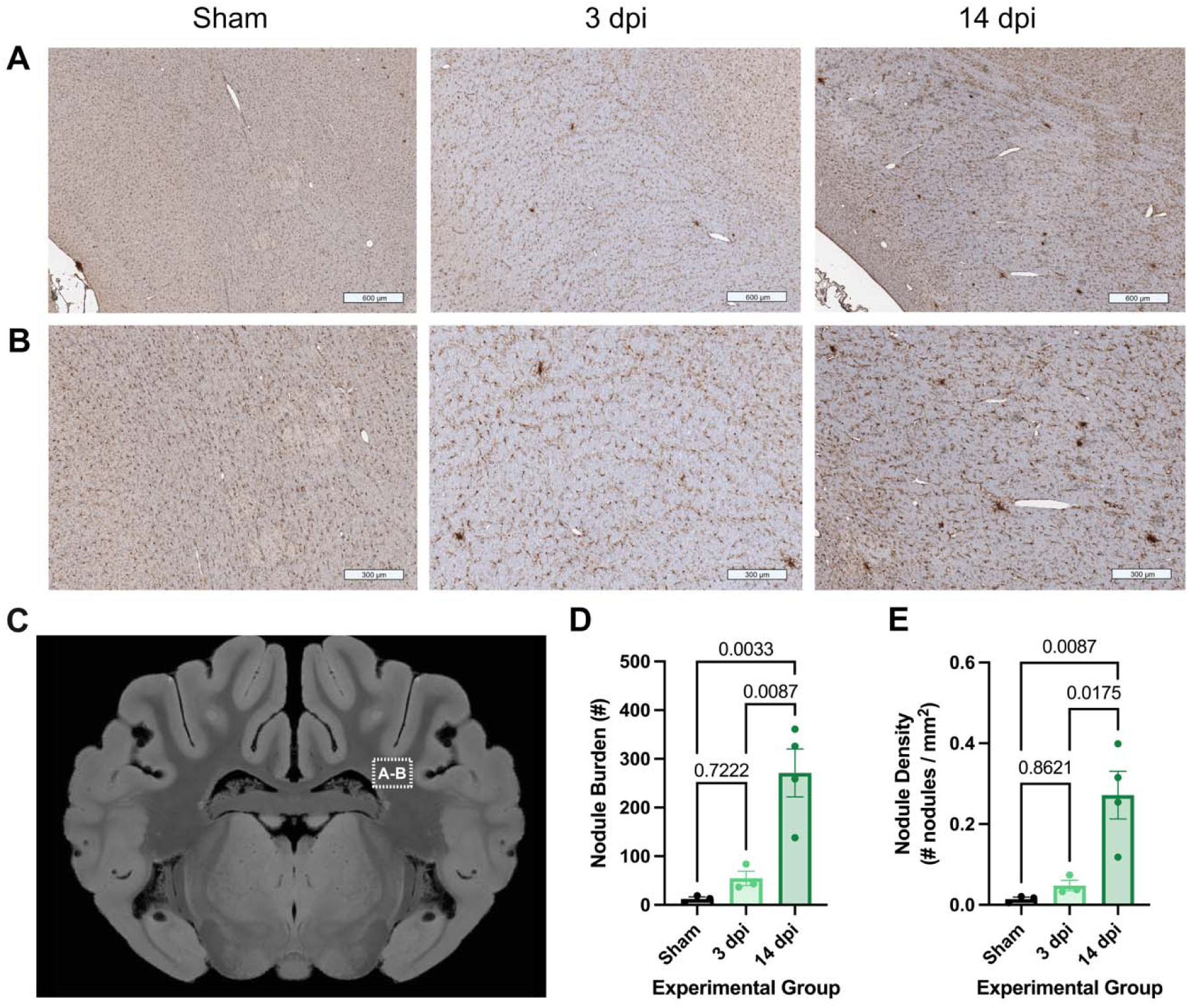
Nodule burden and density increase over time following closed-head rotational TBI. (A–B) Representative Iba1 IHC showing microglia nodules in sham, 3 dpi, and 14 dpi tissue. The upper row shows lower-magnification (scale bars = 600 μm), and the lower row shows the corresponding higher-magnification (scale bars = 300 μm). (**C)** Coronal pig brain image indicating the anatomical region from which representative Iba1-immunostained tissue images in A–B were obtained. (**D)** Quantification of total nodule burden across experimental groups. (**E)** Nodule density across experimental groups. Data are presented as mean ± SEM, with each point representing an individual animal. Statistical significance was assessed using a one-way ANOVA followed by Tukey’s multiple-comparisons test. P values are shown for relevant pairwise comparisons.

### Nodule density varied by region following TBI

Nodules were spread throughout many anatomical regions in the brain. However, to quantify regional distribution of nodule burden and density, we investigated nodules across four regions of interest (ROIs). ROIs were divided into four groups: ROI 1 included cortical gray matter, ROI 2 captured subcortical white matter, corpus callosum, and claustrum, ROI 3 included hippocampal formation and fornix, and ROI 4 consisted of thalamus, basal ganglia, and internal capsule (Fig. 3A-B). Numerous multicellular nodules were observed in ROIs 2, 3, and 4 (Fig. 3).

**Figure 3:**
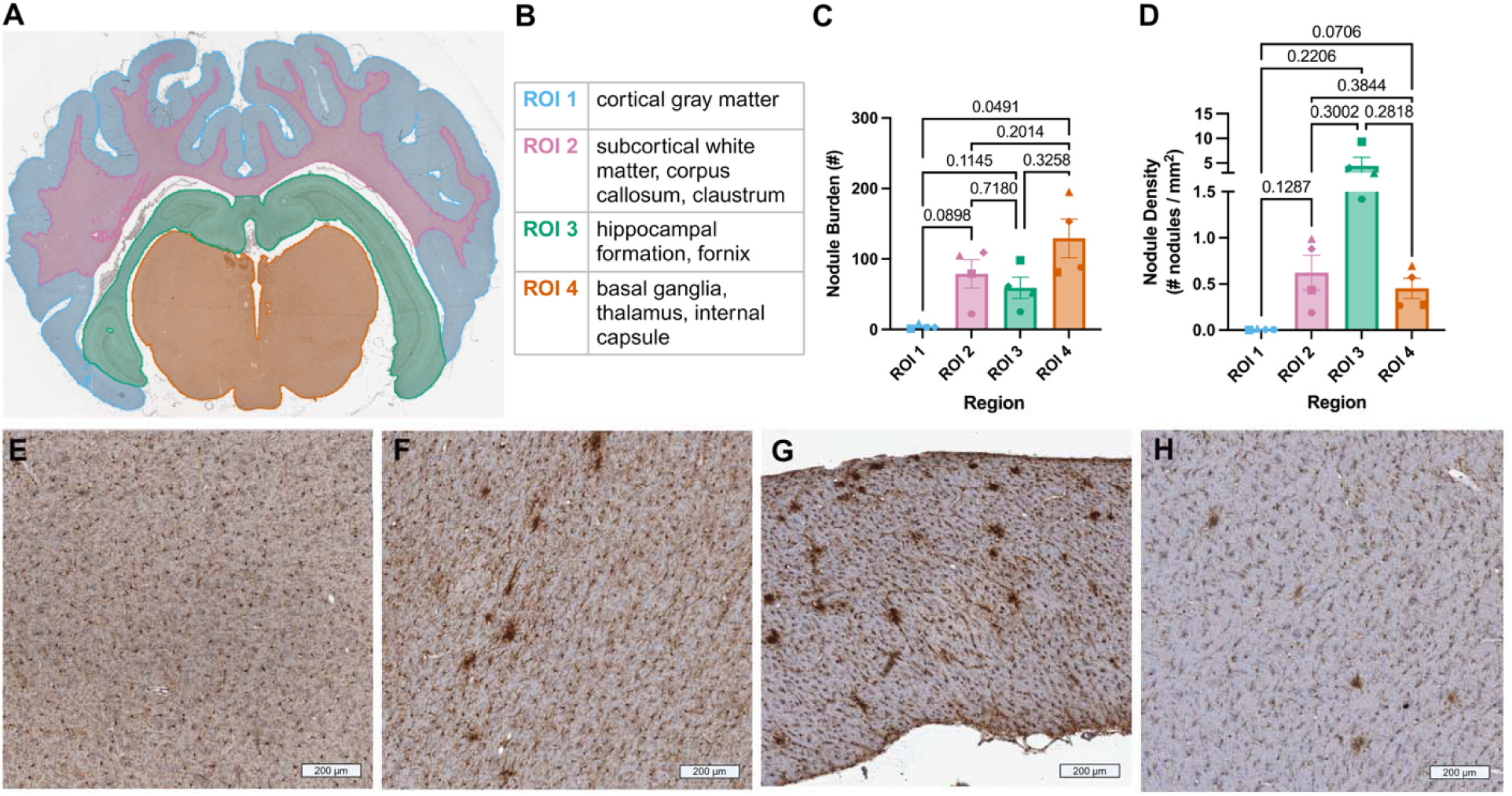
Regional variability in nodule distribution following closed-head rotational TBI. **(A)** Example coronal pig brain section with the four regions of interest (ROIs) used for analysis of nodules. (**B)** Anatomical structures included within each defined ROI. (**C)** Total nodule burden across the four ROIs, 14 dpi. (**D)** Nodule density quantified across the four ROIs, 14 dpi. (**E–H)** Representative Iba1 IHC from (**E**) ROI 1, (**F**) ROI 2, (**G**) ROI 3, and (**H**) ROI 4 in the injured brain (scale bars = 200 μm). Data are presented as mean ± SEM. In **C–D**, each symbol shape represents an individual animal, with the same symbol used to identify a given animal across ROIs. Statistical significance was assessed using a repeated measures one-way ANOVA followed by Geisser-Greenhouse corrections and Tukey’s multiple-comparisons test. P values are shown for relevant pairwise comparisons.

Raw counts demonstrated that ROI 4 contained the greatest nodule burden with a mean 129 27 nodules, followed by ROI 2 with a mean 79 20 nodules, and ROI 3 with a mean 59 15 nodules. In contrast, although the relationship was not statistically significant, ROI 1 had fewer nodules compared to the other ROIs (mean 4 2; Fig. 3C). ROI 4 had a significantly greater nodule burden than ROI 1 (p=0.0491; Fig. 3C). While other ROI comparisons did not reach statistical significance, a trend toward an increased nodule burden was observed between ROI 1 vs. ROI 2 (p=0.0898; Fig. 3C).

Importantly, ROIs were not the same size, so nodule density was calculated by dividing the nodule burden of a region by its respective area. Normalization by ROI area revealed a marked shift in trends. ROI 3 exhibited the greatest nodule density at a mean 4.44 1.71 nodules per mm² (Fig. 3D). ROIs 2 and 4 contained a mean 0.62 0.19 and 0.45 0.11 nodules per mm², respectively (Fig. 3D). ROI 1 consistently showed a minimal burden of 0.01 0.00 nodule per mm² (Fig. 3D). No significant differences in nodule density were detected between ROIs but the comparison between ROI 1 and 4 exhibited a trend towards significance (p=0.0706). Representative images from each region reflected these trends (Fig. 3E-H). Although comparisons between regions lacked statistical significance, these data suggest that there could be anatomical regions that are more susceptible to expressing this unique TBI-induced morphology. For example, ROI 1, containing cortical gray matter, consistently had the lowest nodule burden and nodule density, indicating that nodule formation is relatively uncommon within this region.

### Nodules were enriched within white matter following TBI

To further investigate nodule distribution and the variability observed, we next examined nodule distribution across gray and white matter tissues. ROI 1 exclusively contained gray matter and presented with minimal numbers of nodules. ROI 2 contained mostly white matter structures and presented with a moderate nodule burden and nodule density. ROIs 3 and 4 contained both white and gray matter structures. Numerous multicellular nodules were observed within the white matter spaces of these regions including subcortical white matter, corpus callosum, internal capsule, fornix, and deep white matter (Fig. 4A). In contrast, nodules were less frequently observed in gray matter structures including the gray matter structures of hippocampal formation and gray matter regions of the thalamus and basal ganglia. In line with these observations, nodule burden and density were significantly higher in white matter structures compared to gray matter structures (p=0.0219; p=0.0214; Fig. 4B-C). Most nodules formed primarily along white matter tracts and occasionally at gray-white matter interfaces (Fig. 4D, E, F). Low magnification representative images (Fig. 4E, F) illustrate this anatomical preference, which is also visible in higher magnification. The zoom-ins highlight the nodule distribution within the white matter parenchyma, but not adjacent gray matter spaces (Fig. 4d’, e’, f’). These data suggest that nodules are preferentially enriched throughout the brain in white matter tissues following TBI.

**Figure 4:**
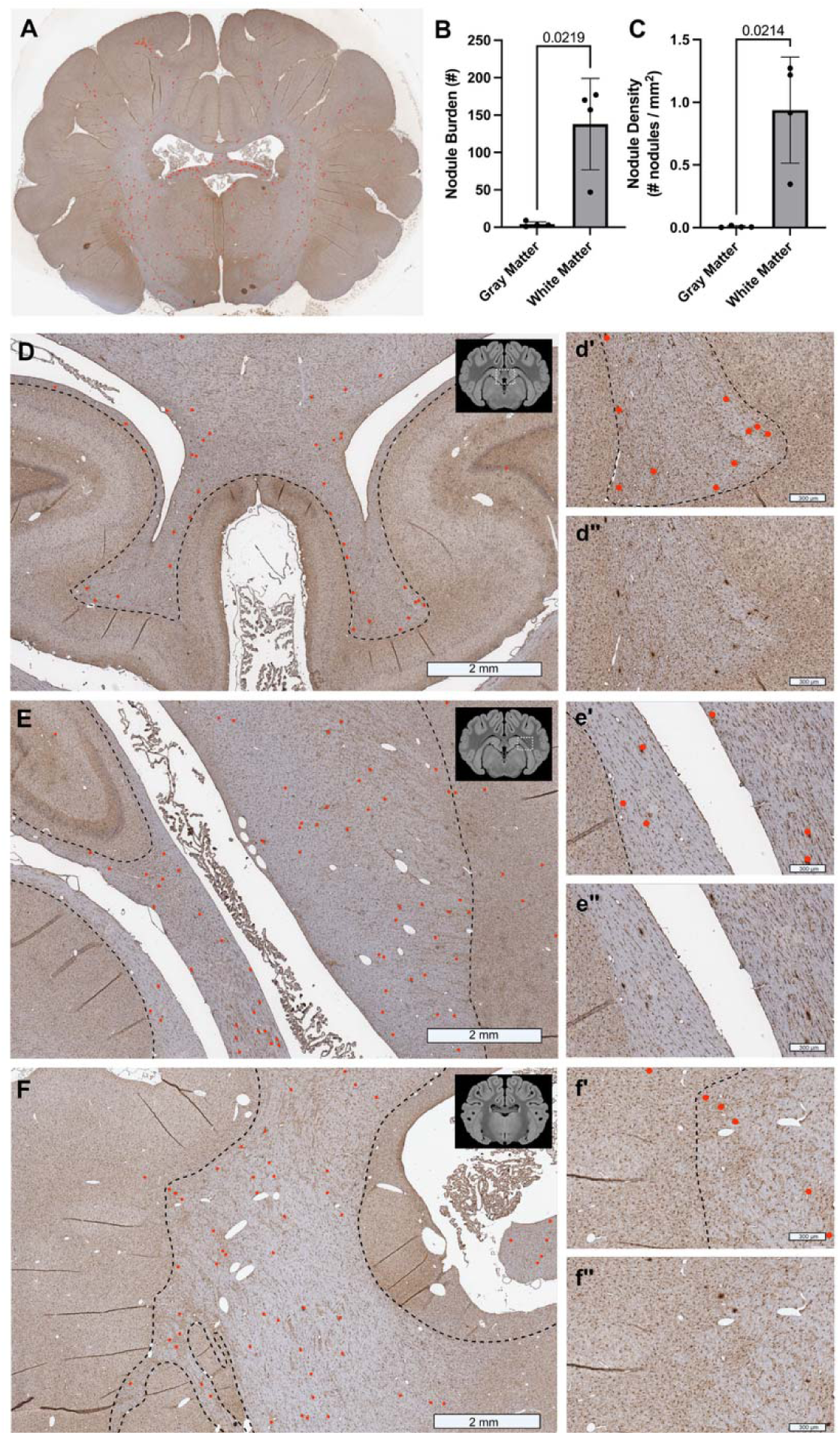
Nodules are predominantly observed in white matter following closed-head rotational TBI in pigs. **(A)** Example 14 dpi pig brain section with nodules labeled by red points. (**B)** Total nodule burden between gray and white matter structures at 14 dpi. (**C)** Nodule density across gray and white matter structures at 14 dpi. (**D–F)** Iba1-immunostained sections illustrating the spatial distribution of nodules across gray and white matter structures (scale bars = 2 mm). Dashed lines separate the boundaries between white and gray matter and red points indicate identified nodules. Brain atlas insets show the anatomical location. (d′, e′, f′) Higher-magnification of areas from (**D-F**), with red points indicating identified nodules and dashed lines indicating white–gray matter boundaries (scale bars = 300 μm). (d″, e″, f″) Corresponding unannotated images. Data are presented as mean ± SEM, with each point representing an individual animal. Statistical significance was assessed using a paired t-test. P values are shown for relevant pairwise comparisons.

### Nodule formation positively correlated with injury velocity

Given that the 14-dpi cohort exhibited the greatest overall nodule burden, we focused our subsequent analyses on this time point across TBI injury severity. Subjects experienced sagittal injuries at either sham, moderate-velocity, or high-velocity (Fig. 5A). Nodule burden after high-velocity TBI was significantly greater than sham (p=0.0032; Supp. Fig. 3C) and moderate velocity injury groups (p=0.0062; Supp. Fig. 3C). Indeed, high-velocity subjects exhibited a 5-fold increase in nodule burden relative to moderate-velocity subjects (Supp. Fig. 3C). Moreover, nodule burden could be expressed as a function of injury velocity, with higher injury velocities corresponding to higher nodule burden (R^2^=0.8383; Fig. 5B). Changes across injury velocity were also conserved in nodule density metrics. High-velocity TBI had significantly elevated nodule density relative to tissue from a moderate-velocity TBI (p=0.0097; Fig. 5C) and sham group (p=0.0062; Fig. 5C).

**Figure 5:**
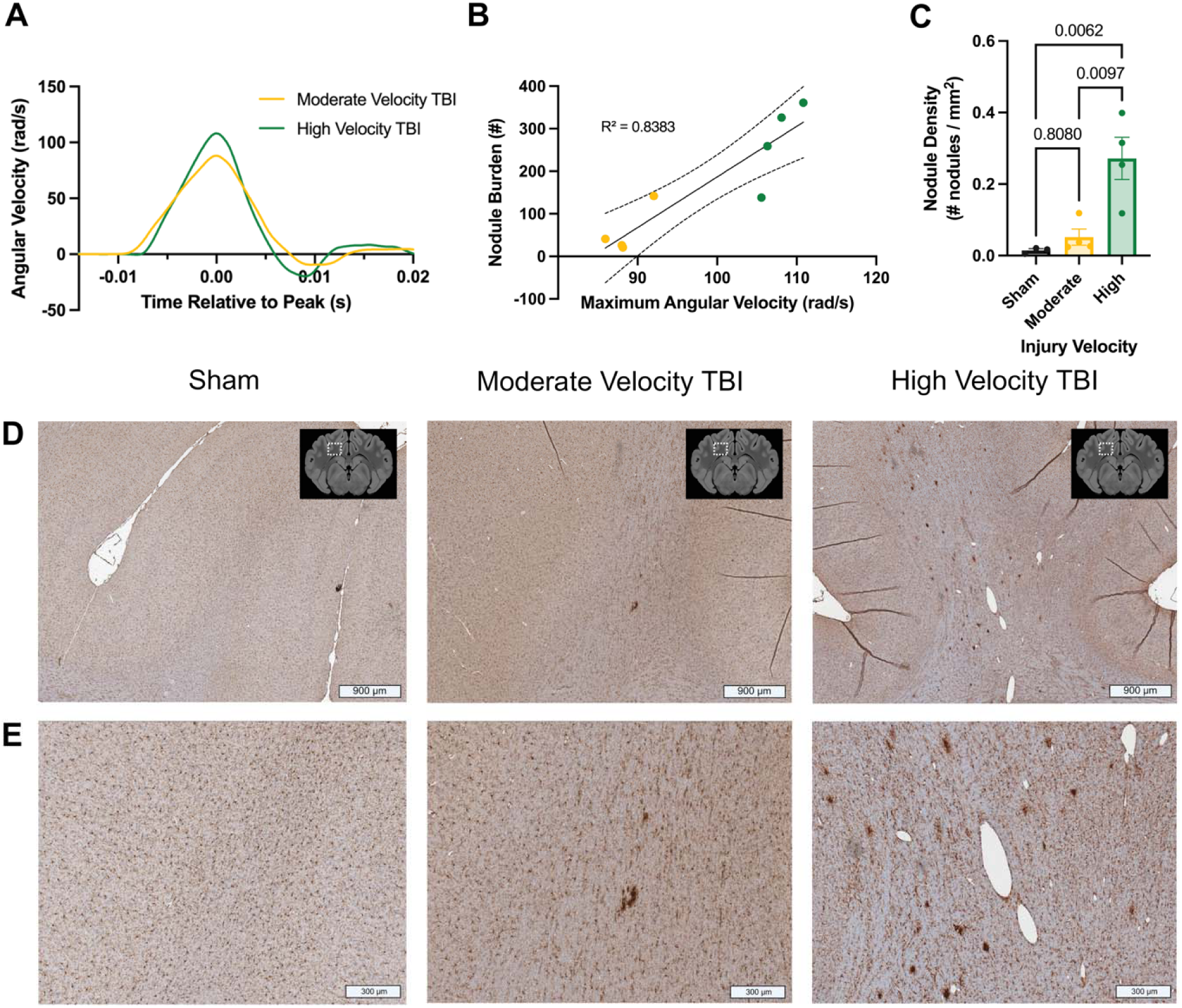
Nodule burden following closed-head TBI is associated with injury severity. **(A)** Example angular velocity traces for moderate and high velocity rotational TBI, which shows differences in peak angular velocity between injury conditions. (**B)** Correlation between maximum angular velocity and total nodule burden in injured animals. (**C)** Nodule density across injury condition. (**D–E)** Iba1-immunostained sections from sham, moderate-velocity TBI, and high-velocity TBI animals. Brain atlas insets indicate the anatomical location of each image. Upper panels (**D**) show lower-magnification (scale bars = 900 μm), and lower panels (**E**) show higher-magnification (scale bars = 300 μm). Data are presented as mean ± SEM, with each point representing an individual animal. Statistical significance was assessed using **(B)** an ordinary least squares estimation and **(C)** a one-way ANOVA followed by Tukey’s multiple-comparisons test. P values are shown for relevant pairwise comparisons.

Although moderate-velocity TBI exhibited a lower overall nodule burden and density, it preserved the regional distribution trends observed in high-velocity TBI group (Supp. Fig. 3A-B). Specifically, ROI 4 contained the greatest nodule burden in moderate-velocity TBI with a mean 27 13 nodules, followed by ROI 3 (15 8 nodules) and ROI 2 (14 7 nodules; Supp. Fig. 3A). ROI 1 consistently showed the lowest burden (3 1 nodules) (Supp. Fig. 3A). Nodule density trends also mirrored those of high-velocity TBI at a smaller scale (Supp. Fig. 3B). ROI 3 exhibited the greatest nodule density at a mean 0.92 0.36 nodules per mm², while ROIs 2 and 4 contained means 0.10 0.05 and 0.08 0.03 nodules per mm², respectively (Supp. Fig. 3B). ROI 1 maintained the lowest nodule density at a mean 0.00 0.00 nodules per mm² (Supp. Fig. 3B). Although comparisons across ROIs lacked statistical significance in the moderate-velocity TBI group, spatial distribution remained consistent with high-velocity TBI trends. Together, these data indicate that nodule formation is modulated by injury velocity with more injurious TBIs correlating to increased nodule burden and density.

## DISCUSSION

Neuroinflammation is a critical determinant of outcomes following TBI. Understanding neuroinflammatory processes that affect humans is critical for basic neurotrauma characterization and could facilitate the development of sensitive diagnostics and translational treatments. To these ends, microglia morphologies including ramified, reactive, amoeboid, rod, hypertrophic, and dystrophic forms have been extensively characterized ^14,48–58^. However, a clinically relevant morphology - multicellular microglial nodules - has been poorly characterized to date. Here, we report that a closed-head model of rotational acceleration diffuse TBI in pigs generates abundant microglia nodules, enriched in white matter spaces at subacute time points.

Importantly, microglia forming into nodules is not unique to pigs but has also been reported in human post-mortem tissue after a TBI ^29–35,59–62^. First described in 1956, only a handful of human post-mortem TBI studies have gone on to characterize their distribution, conditions, and persistence ^32,35,59–61^. Nodules are virtually undetectable at early time points but occur in 75-92% of cases that survive 2 weeks to 2 years post injury ^31,32,34,61,62^. Nodules are present in mild and severe cases after diffuse, impact, or blast TBI in pediatric, adult, and geriatric cases ^30,32,61^. Nodules are abundant in the white matter with particular enrichment in the corpus callosum, internal capsule, and brainstem ^61^.

Nodules are an understudied pathology in TBI. However, nodules are not unique to TBI. They have also been described in multiple sclerosis, stroke, graft-versus-host disease, and Rasmussen encephalitis ^36–39^. Interestingly, nodules have also been observed in post-mortem tissue of patients with Alzheimer’s disease (AD) ^63^. Post-mortem tissue from those with AD and non-demented individuals with high load of AD-related pathologies (i.e., amyloid-beta and hyperphosphorylated tau) had nodules present in the white matter ^63^. In cases with AD pathology, nodules were significantly increased relative to tissue from healthy controls and were associated with the early phases of AD pathophysiology ^63^. Together, these data suggest that immune cell nodules are relevant but not unique to TBI. However, more studies are needed to determine the nodule characteristics and functions that are conserved across diseases.

Given their clinical relevance, we sought to describe TBI-induced nodules in this injury model. Importantly, nodules, which we define as dense aggregations of Iba1+ immune cells, are distinct from glial scarring in their location, initiation, and distribution. Unlike classical glial scarring which typically spans a large area and localizes to the primary injury site or tissue structures in-line with injury forces ^64^, the nodules described here exhibit a distinct distribution profile. Nodules exhibited a pervasive, multifocal punctate distribution, not centralized around one anatomical location. A nodule consists of tightly overlapping cell bodies forming a dense, globular cluster that stand out against normal, ramified microglia. Within nodule-rich white matter, nodules exhibited an approximately even distribution throughout the tissue. Importantly, nodules were surrounded by and were consistently larger than normally appearing ramified microglia. Also, while glial scarring typically appears at acute time points ^65–68^, nodules have a delayed onset, with robust numbers occurring in the subacute phase (14 dpi). Finally, nodules appear evenly distributed throughout a given region (e.g., fornix, claustrum, and corpus callosum) but they have region-to-region variability with some regions exhibiting high nodule burden (e.g., fornix) and other regions exhibiting almost no nodule burden (e.g., cortical gray matter). Together, nodules are distinct from glial scarring because of their distribution pattern, morphology, temporal profile, and tissue-structure specificity.

Similar multicellular myeloid cell nodules are not unique to humans and pigs but have been described in other animal models as well. For instance, Iba1+ nodules have been reported within the subcortical white matter of the injured hemisphere following a mild controlled cortical impact (CCI) in ferrets ^69^. In this ferret model, Iba1+ nodules developed in subcortical white matter at subacute time points after TBI. Nodule burden peaked 4 weeks post injury and remained at elevated levels out to 16 weeks post injury, the latest time point considered ^69^. Likewise, Iba1+ nodules have been noted in rodent TBI models as well. For example, nodules have been observed in the optic nerve and optic tract of mice after a closed-head weight drop model ^70,71^. However, nodule burden is quite distinct across these models. Nodule incidence in rodents is rare and inconsistently described across injury models ^53,54,58,70–73^. Nodule burden in ferrets peaked approximately 15 nodules per coronal cross-section. In contrast, nodule burden in this pig injury model numbers in the hundreds. This increase in nodule burden exceeds the 10-fold difference in brain mass ^74^. Considering that rodent brains are lissencephalic with lower proportion of white matter, these data suggest that nodules form independently of injury model but do occur with larger numbers in brains with larger mass and gyrencephalic architecture. However, why nodules are more abundant in ferret, pig, and human brains has yet to be determined.

Like ferret and human postmortem observations, our porcine data showed that nodule burden following TBI was primarily concentrated within white matter structures. While damage during closed-head diffuse TBI can occur in both gray and white matter tissues, it is widely appreciated that diffuse axonal injury in white matter tracts is a hallmark pathology of closed-head diffuse TBI ^75,76^. Thus, enrichment of nodules in white matter could be caused by white matter-specific debris, diffuse axonal injury, or localized environmental signals. However, sparse populations of nodules are still present in the gray matter. Future studies should investigate the stimuli for nodules and whether the transcriptomic profiles or functional states of gray matter nodules mirror those found in the white matter.

While our current experimental design incorporates a sham control, an acute time point (3 dpi), and a subacute time point (14 dpi), a substantial increase in nodule burden is observed between 3 and 14 dpi. Consequently, the precise temporal dynamics and the exact window of initial nodule formation remain unknown. Notably, the preservation of regional distribution patterns across varying injury severities suggests that the underlying anatomical vulnerability is consistent, with severity dictating only the magnitude of the response. However, because our observations are limited to a subacute window, it remains unclear whether the burden observed at 14 dpi represents a pathological peak of the given injury level or if these structures continue to accumulate, plateau, or undergo resolution over time. Longitudinal studies mapping the full trajectory from acute formation to chronic survival are essential to determine whether these nodules persist, amplify, or resolve. These data may also yield insights to understanding whether nodules are adaptive or maladaptive responses to TBI and define potential windows for therapeutic modulation.

Although this study provides valuable insights into understudied myeloid cell nodules in a porcine TBI model, several limitations should be considered when interpreting the results. First, the study includes only female subjects. Previous human literature has included both male and female subjects, but did not note any sex-dependent differences ^29,35,61^. However, male pigs should be included in future analyses to control for possible sex-dependent differences in nodule formation and dynamics. Second, the small sample size needs to be considered when interpreting the statistical analyses to account for potential Type II errors. However, given the trends towards increased nodule burden or density, larger sample size may help further clarify this relationship. Lastly, while previous literature often classifies these structures as “microglial clusters” or “activated microglia,” standard markers like Iba1, HLA-DR, and CD68 stain for both resident microglia and infiltrating peripheral immune cells ^29,63,77–79^. In-depth analysis of their precise cellular composition will be necessary to establish whether these nodules are strictly microglial or composed of mixed myeloid populations. Additionally, it is unknown whether this morphology results from the aggregation of individual, adjacent immune cells or from their fusion into multinucleated cells.

In summary, this study establishes myeloid cell nodules as a robust and reproducible neuroinflammatory endophenotype following closed-head diffuse TBI in pigs. Nodule burden increased as a function of both time and injury severity and was enriched within white matter regions. This immune cell endophenotype mimics presentations in human tissue and although the precise functions of nodules remain unknown, their delayed development and anatomical distribution suggest they contribute to white matter tissue dynamics. Future studies investigating the persistence, composition, functions, phenotypes, and associations of nodules may clarify their contributions following trauma.

## CONCLUSION

Neuroinflammation impacts chronic neurological dysfunction and long-term recovery following TBI. Although multicellular myeloid cell nodules have been described in human tissue, they remain understudied in preclinical models. Here, we demonstrated that a closed-head diffuse TBI model in pigs reliably replicates this human pathology. Nodule burden increased over time and with injury severity, showing a strong preference for white matter regions with negligible presence in the cortical gray matter. These findings establish myeloid cell nodules as a distinct and quantifiable response to diffuse brain injury and provide a foundation for future investigations into their cellular composition, biological function, and contribution to long-term tissue remodeling after TBI. Understanding the role of these nodules may reveal novel mechanisms linking TBI to persistent changes in brain health.

## TRANSPARENCY, RIGOR, AND REPRODUCIBILITY SUMMARY

All animal care and procedures were performed in accordance with the IACUC and following ARRIVE guidelines at the University of Pennsylvania. Animals were randomly assigned to groups. Nodule analysis was not the primary goal of animal enrollment in this cohort, instead nodule observations were systematically assessed in this characterization study. However, animal enrollment of n=3-4 subjects per condition in this study is in line with previous large-animal manuscripts. All experimental analyses were performed by experimenters blinded to the experimental groups. All nodules within each ROI were labeled and quantified. Non-parametric statistical tests were utilized to assess differences between animal conditions and regions since equal variance assumptions of parametric tests were violated. All data presented in the article were independently reviewed and verified by multiple authors. Data supporting the findings of this study are publicly available at Open Data Commons for Traumatic Brain Injury (ODC-TBI), DOI: 10.34945/F5XP4C.

## AUTHORSHIP CONFIRMATION/CONTRIBUTION STATEMENT

Project conception was completed by KLW. SS, AT, CCL, NM, and AA conducted image quantification, data processing, statistical analyses, and results interpretation. KDB and KLW acquired data through animal enrollment, injury processes, tissue collection, staining, and imaging. MLB, DKC, DFM, and KLW provided funding, guidance, and supervision of the project. SS and KLW wrote and edited the manuscript. All authors read, edited, and approved the final manuscript.

## Supplement

**Supplemental Figure 1:**
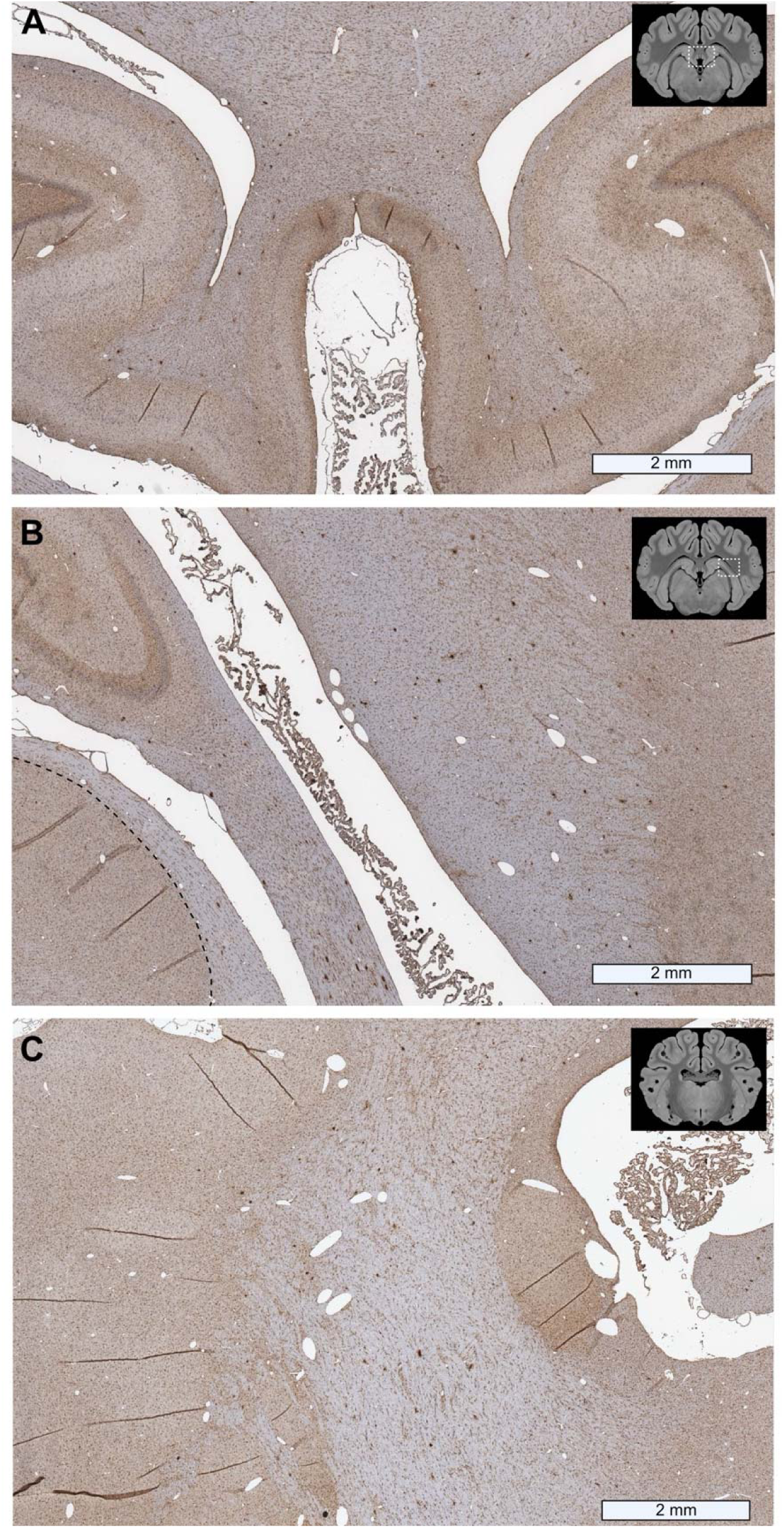
**A–C** Iba1-immunostained sections illustrating the spatial distribution of nodules across gray and white matter structures (scale bars = 2 mm). Corresponding unannotated images of Fig. 4D-F. Brain atlas insets show the anatomical location.

**Supplemental Figure 2:**
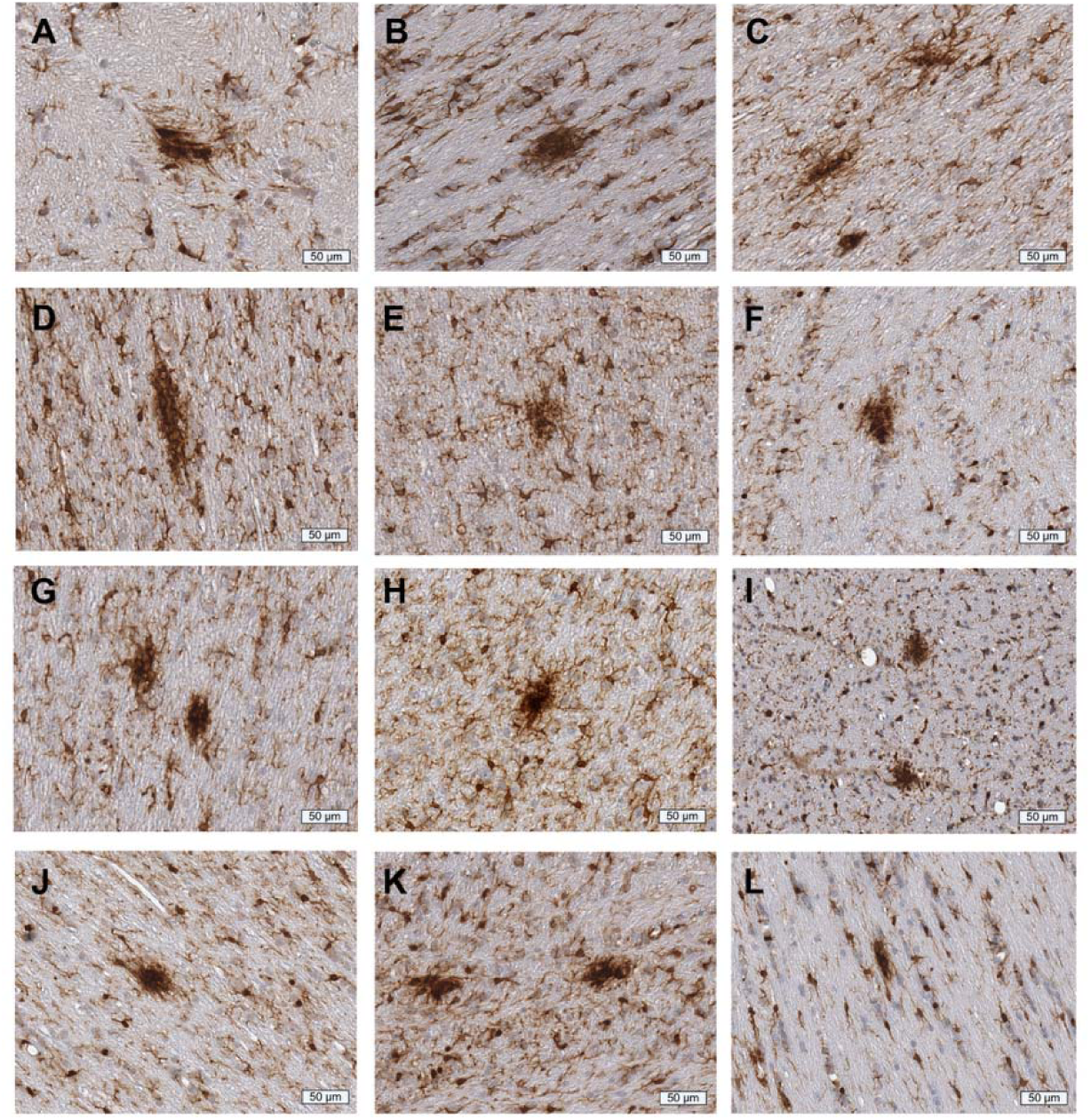
**(A–L)** Representative high-magnification images of nodules found throughout the tissue across brain regions, injury severity, and survival timepoints demonstrating various characteristics (scale bars = 50 μm).

**Supplemental Figure 3:**
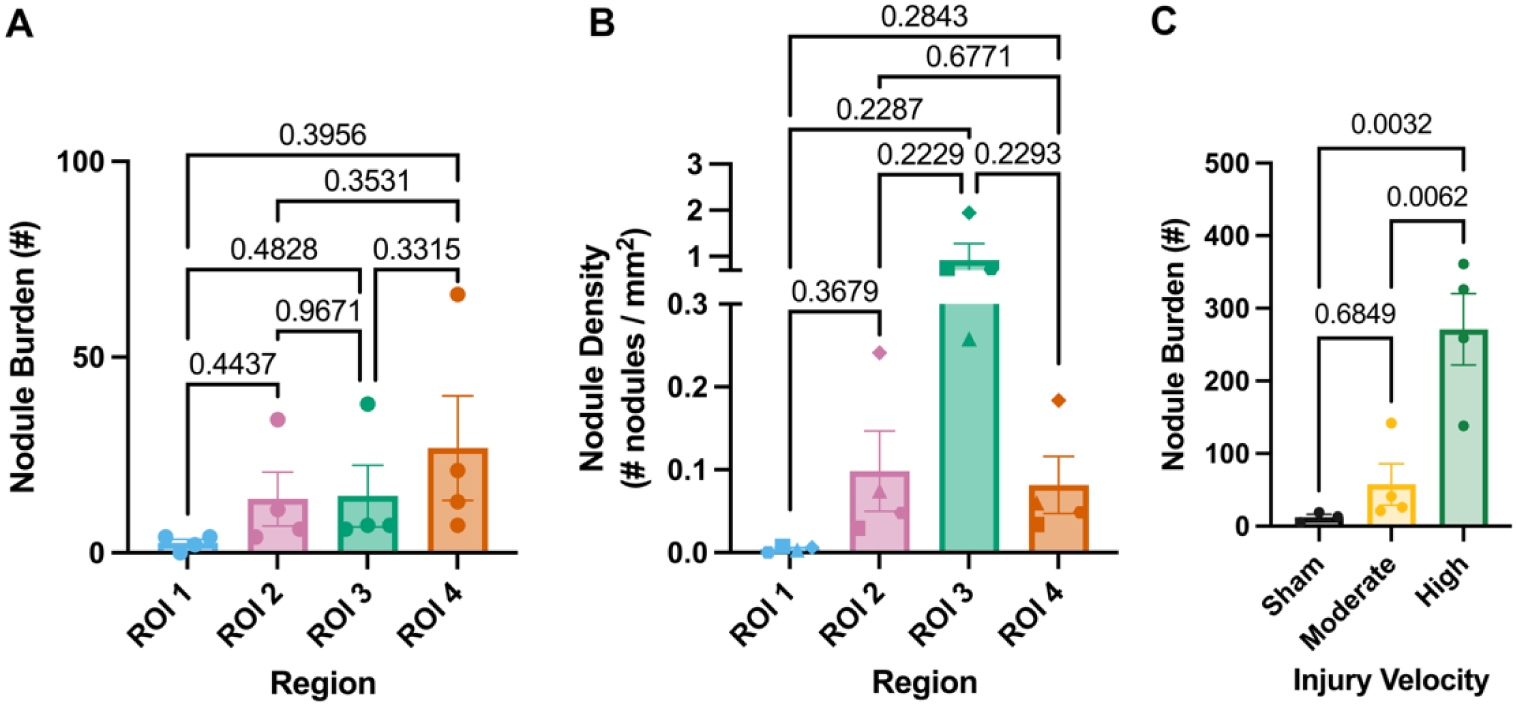
**(A)** Nodule burden across ROIs after a moderate velocity TBI. (**B)** Nodule density across ROIs after a moderate velocity TBI. **(C)** Nodule burden across injury condition. Data are presented as mean ± SEM, with each point representing an individual animal, with the same symbol used to identify a given animal across ROIs. Statistical significance was assessed using **(A-B)** a repeated measures one-way ANOVA followed by Geisser-Greenhouse corrections and Tukey’s multiple-comparisons test and **(C)** a one-way ANOVA followed by Tukey’s multiple-comparisons test. P values are shown for relevant pairwise comparisons.

## DECLARATIONS

### ETHICAL CONSIDERATIONS

Not applicable.

### CONSENT TO PARTICIPATE

Not applicable.

### CONSENT FOR PUBLICATION

Not applicable.

## Acknowledgements

Not applicable.

